# Better tired than lost: turtle ant trail networks favor coherence over short edges

**DOI:** 10.1101/714410

**Authors:** Arjun Chandrasekhar, James A. R. Marshall, Cortnea Austin, Saket Navlakha, Deborah M. Gordon

## Abstract

Creating a routing backbone is a fundamental problem in both biology and engineering. The routing backbone of the trail networks of arboreal turtle ants (*Cephalotes goniodontus*) connects many nests and food sources using trail pheromone deposited by ants as they walk. Unlike species that forage on the ground, the trail networks of arboreal ants are constrained by the vegetation. We examined what objectives the trail networks meet by comparing the observed ant trail networks with networks of random, hypothetical trail networks in the same surrounding vegetation and with trails optimized for four objectives: minimizing path length, minimizing average edge length, minimizing number of nodes, and minimizing opportunities to get lost. The ants’ trails minimized path length by minimizing the number of nodes traversed rather than choosing short edges. In addition, the ants’ trails reduced the opportunity for ants to get lost at each node, favoring nodes with 3D configurations most likely to be reinforced by pheromone. Thus, rather than finding the shortest edges, turtle ant trail networks take advantage of natural variation in the environment to favor coherence, keeping the ants together on the trails.

**Author Summary:** We investigated the trail networks of arboreal turtle ants in the canopy of the tropical forest, to ask what characterizes the colony’s choice of foraging paths within the vegetation. We monitored day to day changes in the junctions and edges of trail networks of colonies in the dry forest of western Mexico. We compared the paths used by the ants to simulated random paths in the surrounding vegetation. We found that the paths of turtle ants prioritize coherence, keeping ants together on the trail, over minimizing the average edge length. The choice of paths reduces the number of junctions in the trail where ants could get lost, and favors junctions with a physical configuration that makes it likely that successive ants will reinforce the same path. Our work suggests that design principles that emphasize keeping information flow constrained to streamlined, coherent trails may be useful in human-designed distributed routing and transport networks or robot swarms.

## Introduction

Many engineered systems rely on a backbone routing network, whose goal is to ensure that any two entities or devices on the network can communicate through some path [1, 2, 3]. Some biological systems, such as neural arbors [4], plant arbors [5], and slime molds [6], also use routing networks to transmit information and nutrients. Effective design of routing networks depends on the physical environment, because variation in the environment can affect the accuracy and rate of communication in both engineered [7, 8, 9] and evolved natural networks [10, 11, 12, 13]. The environment influences how the system chooses search strategies, prioritizes competing objectives, and coordinates its local decisions. For example, wireless networks operating in difficult to reach environments may use different routing strategies to minimize energy consumption of devices [7]; similarly, in bacterial navigation, chemicals appearing as localized pulses in the environment can affect gradient sensing and movement patterns [12].

The 14,000 species of ants have evolved diverse distributed routing algorithms to search for, obtain, and distribute resources [14, 15] in diverse environments [16, 17, 18, 19, 20]. Models of engineered routing networks inspired by ants often emphasize the goal of minimizing the distance traveled. Ant colony optimization (ACO), first proposed in 1991, loosely mimics ant behavior to solve combinatorial optimization problems, such as the traveling salesman [21, 22, 23] and other routing problems [24]. In ACO, individual ants each use a heuristic to construct candidate solutions, and then use pheromone to lead other ants towards better solutions. Recent advances improve ACO with techniques such as local search [25], cunning ants [26], and iterated ants [27].

A fascinating recent area of biological research examines the goals met by the trail networks of ants [20]. Studies of species that forage on a continuous 2D surface [18], including Pharaoh’s ants [28], Argentine ants [17, 29, 30], leaf-cutter ants [16], army ants [31, 32], red wood ants [33], and meat ants [34, 35], show that ants use local chemical interactions to form trails [36, 37, 31, 38], regulate traffic flow [39], search collectively [40], and form living bridges [41].

There are many objectives that an ant colony’s trail network might meet, including minimizing costs in time and energy by reducing the distance traveled, keeping the ants together to form a coherent trail, being resilient to rupture, and searching effectively [42]. Ant species that forage and form trails on the ground have few constraints on trail geometry because their trails can form nodes, junctions where ants choose among more than one direction or edge, and edges anywhere on the 2D plane. One mathematical model for trail network objectives draws on the classic Euclidean Steiner tree problem, in which the goal is to connect terminal nodes, such as nests and food sources, in the plane with minimal total length, measured using Euclidean distance [43]. For example, Argentine ants observed in a lab setting form foraging trails that mimic approximate Steiner trees [17]. However, other studies of polydomous species in the field indicate that minimizing the distance traveled may not be the only objective that ant trail networks attempt to optimize. Many polydomous species construct trails that increase robustness through redundant connections, which would not be predicted by the Steiner tree model [42]. Red wood ants manage trade-offs between minimizing total trail length and minimizing the time needed to travel between the nest and each food source [44], as well as trade-offs between nest centrality and robustness [45]. Meat ants form trail networks that link nests and trees as nodes, and their choices of which nodes are linked, as well as the direction and length of trails, suggest that robustness to the loss of a node is as important as minimizing the distance traveled [34, 35]. When given the choice of two paths of equal length, black garden ants prefer straight paths to curved paths [46].

While many ground-living species can create nodes and edges at any location in a 2D plane, studies of many ant species show that physical characteristics of nodes in trails, such as the angle of bifurcation [47, 30], influence the structure of trail networks. Army ants sometimes link their bodies to form a bridge across gaps that shortens the path, but do not always make bridges because it reduces the number of available foragers [48].

The arboreal turtle ant (*Cephalotes goniodontus*) nests and forages in the tree canopy of tropical forests [49]. A turtle ant colony creates a trail network in the vegetation that connects several nests, providing a routing backbone that must be maintained to allow resources to be distributed throughout the colony [50, 51]. Trail networks appear to be maintained by trail pheromone, which the ants lay as they walk along. In previous work, we used observational data to parameterize a model for how ants select a trajectory through a node to the next edge, based on the rate at which volatile pheromone is deposited and decays [52]. Simulation results were consistent with field observations [50] indicating that at every node, there is a small probability that an ant may leave the trail to explore, by taking an edge that is not the one most strongly reinforced by pheromone [52]. While a colony’s trail network is often similar from day to day [50], trails tend to change at least a few nodes on the timescale of days, while nests change on the timescale of weeks [50, 51]. One source of day to day changes is the formation of new trails to temporary food sources, such as nectar, fungi, and probably pollen, which are eventually depleted. This species nests in burrows in rotten wood that were created by larval beetles [53]. Colonies sometimes add a new nest [54] or lose a nest when the branch a nest is in breaks off [51]. Day to day changes in trails sometimes include small alternative paths, or loops, of which one is eventually chosen. The colony also repairs the backbone in response to frequent changes in the vegetation caused by plant growth and by ruptures made by wind or passing animals [51].

## Results

### Candidate objectives

We used data from field observations of the trail networks of turtle ants to test what objective functions these trails may be optimizing. We compared the observed networks with simulated random networks, to determine how well the observed networks meet four objectives.

1. Minimizing the average length of the edges in the trail network. This objective, which is equivalent to minimizing the total trail length for a fixed number of edges, contributes to reducing the costs in energy and time of travelling through the trail network. The distance between nodes is not uniform, so the total length of a path is not equivalent to the number of nodes traversed, as is often assumed in various optimization algorithms [55, 24, 56, 57], including our previous work on turtle ants [52], because in the vegetation, the lengths of edges vary; the distance between one node and another ranges over 100-fold, from less than a centimeter to more than a meter (Table 1).
2. Minimizing the total number of nodes, which promotes the maintenance of a coherent trail by reducing opportunities for ants to get lost. The number of nodes was a count of all the nodes traversed in the trail network. Because ants explore edges not reinforced by pheromone, each node presents an opportunity for ants to get lost, and lost ants may lay pheromone trail that could lead other ants astray. Each node is also an opportunity to intersect with the network of a competing colony [58] in the same vegetation.
3. Maximizing the probability that successive ants take the same trajectory through a node. This objective contributes to the creation and maintenance of a coherent trail. The transition from one edge through a node to another edge is reinforced only when successive ants deposit pheromone along exactly the same trajectory through the node (Fig 1); the ants have short antennae that detect pheromone only locally. Nodes are variable 3D structures, ranging from a simple fork created by a branch in a plant, to a cluster of entwined vines and branches from different plants. We used a categorical index, based on the physical characteristics of the vegetation at each node, to estimate how many trajectories an ant could take from one edge through a node to another edge, and thus how likely a node is to be reinforced (Fig 1). Each transition through a node in the vegetation (e.g., edge *u* → *v* through node *v* to edge *v* → *w*), both along the trail used by the ants and in the surrounding vegetation, was assigned a value of a transition index. Criteria for assigning values of the transition index are listed in the legend for Fig 1. These criteria were not based on any assessment of the behavior of the ants, and the same procedure to evaluate plant characteristics was used for both the nodes along the ants’ trails and the nodes in the surrounding vegetation. Values of the transition index ranged from 1, assigned when the node and both edges were all on the same plant, so successive ants were most likely to take and reinforce the same trajectory through the node (Fig 1A), to 4, assigned when a gap between stems or interference from the wind made it unlikely that successive ants could take the same path (Fig 1D). Different trajectories from one edge to another along a node may be assigned different transition indices. This index is not a measure of the biomechanical difficulty of a particular transition; ants do not seem deterred by features such as the 3D angle of two edges at a junction, except in very extreme cases [51]. We used the average transition index of all transitions in the trail network to test whether the ants’ choice of nodes reflected our assessment of the probabilities that certain trajectories would be reinforced. This extends our previous work [51, 52] which did not take into account the physical variation among nodes, and did not examine how trails are selected from among the many available alternative networks.
4. Minimizing the path length (i.e., the total length of all edges in the network), which measures the total distance traveled. This is a common measure used to assess the quality of trails for many species [17, 44]. While minimizing the average edge length (first objective) or the number of nodes (second objective) does not guarantee that the path length is minimized, in practice, it is common for at least one of these two objectives to be correlated with path length. Thus, if ants minimize path length, we can then examine whether they do so by minimizing number of nodes, average edge length, or both.

**Fig. 1:**
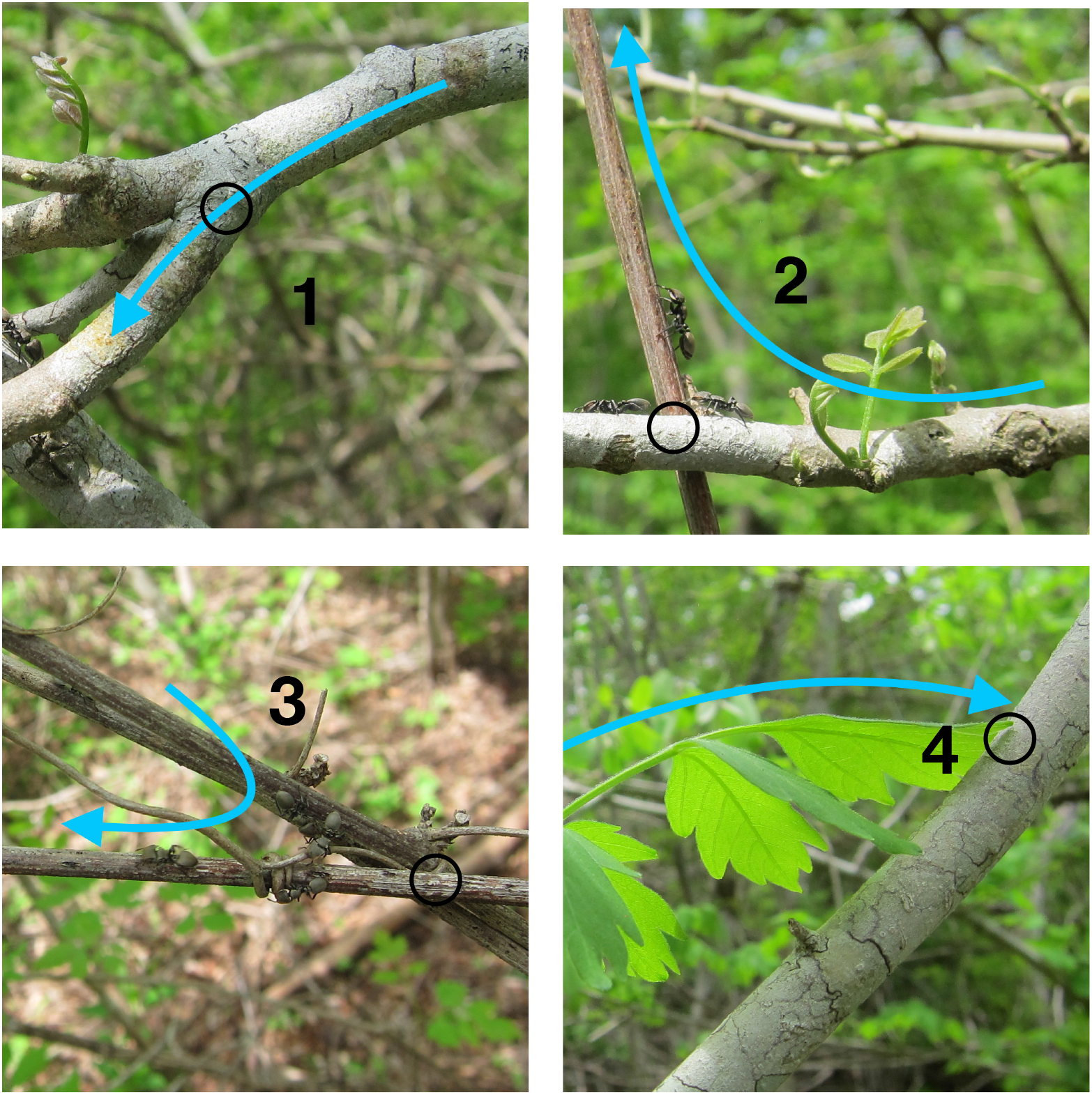
Transition indices. The photos show examples of nodes of each transition index (TI). The open black circle shows the node traversed. The blue arrow shows the transition, leading from the edge along which ants enter the node, through the node, toward the edge along which they exit. Each transition, from an edge to a node to another edge, was assigned a transition index with a value between 1 and 4. A low transition index corresponds to a transition in which the configuration of plants offers only one available trajectory, and thus likely to be rapidly reinforced by successive ants traveling along the same trajectory. TI–1 (upper left): a node linking two edges on the same plant; in the example shown, all ants are likely to walk the same way across the top of the branch. TI–2 (upper right): a node that links one plant to another in which the vegetation offers one very probable trajectory to traverse the node, likely to be used by successive ants; in the example shown, most ants are likely to climb up the brown vine from the position shown by the ant approaching the junction from the left. TI–3 (lower left): a node that links one plant to another plant with more than one possible trajectory through the node; in the example shown, ants on the upper vine can reach the lower one either directly or by following the smaller vine that ants in the photo are using. TI–4 (lower right): a node that links one plant to another with many possible trajectories that are often changed by conditions; in the example shown, wind can easily move the leaf so that different ants reach the junction between the leaf and branch, at different places.

**Table 1:** Comparison of random, optimized, and observed networks. A) Similarity of random, optimized, and observed networks measured as mean percentile (Methods). A percentile of 50 (dashed line) indicates that the observed or optimized network optimized the objective to the same extent as the average random network; the lower the percentile, the better the network optimized the objective compared to random networks. Error bars show standard errors of the mean of 36 non-independent networks. Asterisks represent p < 0.001, Conover test. B) A swarmplot of the percentile of each observed network, as well as a sample of random networks. Gray circles represent the performances of random networks, and colored x’s represent the performances of observed trail networks. Values below 50 indicate performing better than the median random network for that objective, and values above 50 represent performing worse. Asterisks represent p < 0.001, Conover test.

|  | Edge Length (cm) |  | Total Nodes |  | Transition Index |  | Connectivity |
| --- | --- | --- | --- | --- | --- | --- | --- |
|  | Avail | Used | Avail | Used | Avail | Used |  |
| **Tejon 189** | 10698 | 20.64 ± 6.17 | 217 | 43.50 ± 9.43 | 1.60 | 1.31 ± 0.06 | 3.75 ± 0.23 |
| **Tejon 446** | 7625 | 13.48 ± 0.71 | 196 | 35.91 ± 4.06 | 1.69 | 1.44 ± 0.13 | 4.00 ± 0.24 |
| **Tejon 500** | 7341 | 26.20 ± 2.43 | 147 | 25.20 ± 6.94 | 1.88 | 1.60 ± 0.12 | 2.46. ± 0.34 |
| **Turtle Hill 460** | 14696 | 40.00 ± 9.39 | 202 | 54.20 ± 18.23 | 1.47 | 1.44 ± 0.13 | 4.50 ± 0.92 |

### Mapping and modeling turtle ant trail networks

To examine what objectives are met by the ants’ choice of paths within the vegetation (Fig 2A), we mapped the trail networks that connected the nests and naturally occurring, ephemeral food sources of three colonies (Tejon 189, Tejon 446, and Turtle Hill 460), for 10–15 days over the course of 6 weeks in 2018, and one colony (Tejon 500) for 5 days during 1 week in 2019, in a tropical dry forest at La Estación Biológica de Chamela in Jalisco, Mexico.

**Fig. 2:**
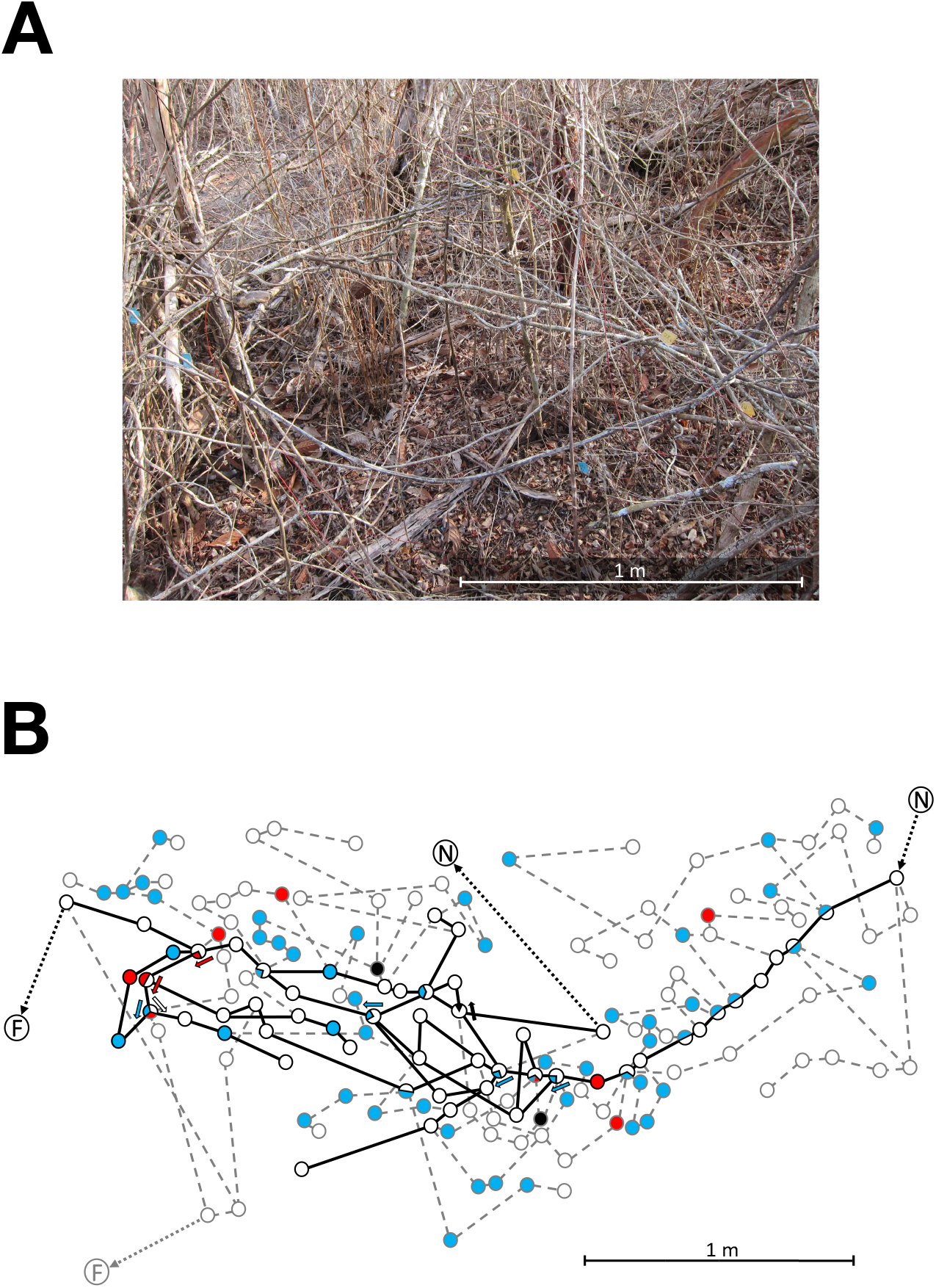
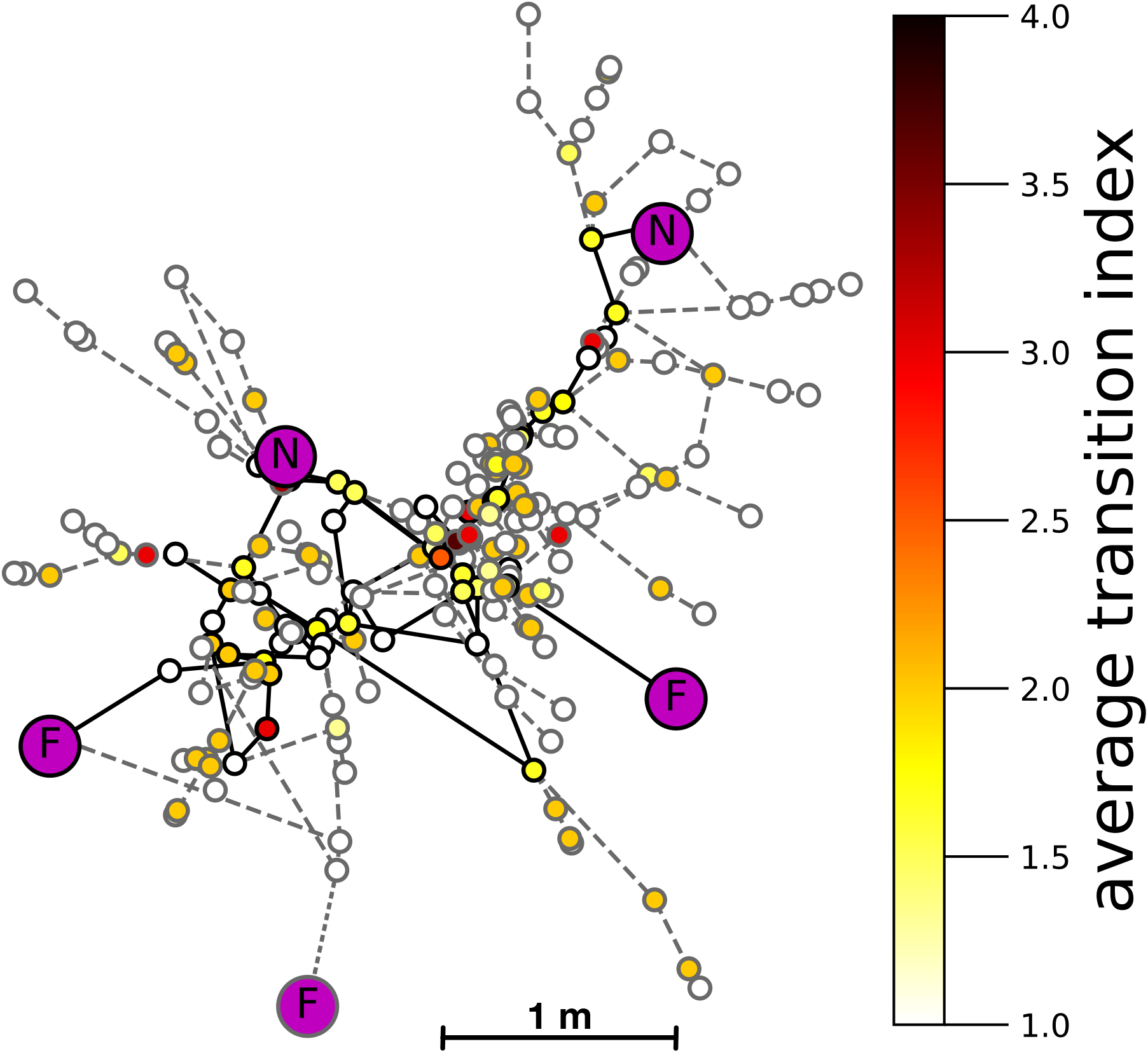
Used and available networks in the vegetation. A) Vegetation in which the trail network mapped in B was made, photographed in the dry season before the branches have leaves. B) Illustration of part of the trail network and network of surrounding vegetation for Tejon 189 on day 9. The figure shows 166 of the 217 nodes mapped in the surrounding vegetation. For many nodes, not all edges are shown to simplify the illustration. Edge lengths are scaled to measured distance, but actual location is not represented here. N represents a nest, F represents a food source. Circles represent nodes. Solid lines represent edges used on that day; dashed lines represent edges not used that day (Methods). The color of a node represents the transition index (TI) from the preceding edge to the following one: TI–1, open circles; TI–2, blue; TI–3, red; TI–4, black. At a node where there is a choice of more than one edge in the indicated direction, so that there could be more than one transition taken through a given node, a TI was assigned to each possible transition. For such nodes with more than one transition index, the TI is represented graphically with a pie chart, and arrows show which transition has the TI represented by the arrow’s color. C) Map of all 217 nodes for Tejon 189 on day 9. Symbols as in B. Not all edges are shown. The color of the node corresponds to the average transition index of transitions through the node. Darker colors represent higher average transition indices. Purple nodes designate nests (N) and food sources (F).

On each day, in each colony, we mapped the trail network, i.e., all paths taken by the ants that had a rate of flow ranging from 10 to 40 ants per 15 mins [51]. We identified each node or junction in the vegetation where an ant had a choice among more than one edge in the direction it was traveling. We measured the length of each edge, and assigned a transition index to each transition, from one edge through a node to another edge, by estimating how likely successive ants would be to take the same trajectory through the node and thus reinforce it with pheromone. To evaluate how the ants choose nodes and edges from the options provided by the surrounding vegetation, we also mapped most of the nodes, measured most of the edges, and assigned transition indices, for all possible paths up to five nodes away from each node used by the ants. A trail network on one day and some of the surrounding network of vegetation is illustrated in Fig 2B–C. From one day to the next, terminals in trail networks may be added or discarded, as new nests are found and food sources are depleted, and nodes within paths are added or deleted [51] (Fig. S1). See Methods for further details.

We modeled the network of vegetation as a directed, weighted graph, *G* = (*V_G_*, *E_G_*), where each junction forms a node, and edges represent stems or branches that connect one node to another. Edge weights correspond to physical length. We modeled transition indices by converting *G* into its corresponding line graph (Methods). The nodes corresponding to the nests and food sources were designated as terminals. There were sometimes day to day changes in which terminals the ants’ paths included in the network.

We compared the extent to which each observed network on a given day, connecting the terminals used on that day, optimized each of the 4 objectives, relative to a set of 100,000 random networks that connected the same terminals used in the observed networks. We compared the observed networks with random networks based on two null models (Methods): For the first null model, the random network generated on day *t* modifies the random network generated on day *t* − 1 by removing paths to terminals not used on day *t* and adding new random paths to terminals used on day *t* but not used on day *t* − 1. This reflects the observation that trail networks are similar from day to day [51]. The second method generates a new random network from scratch each day, so each day is independent of the others.

To compare observed and random networks for each objective, we used a percentile measure (Methods). The percentile measure evaluates how much better the observed network does at optimizing each objective than random networks. The lower the percentile, the better the observed network optimized the objective, and a percentile of 50% means the observed network attains the same value as the median random network.

To determine which of the four objectives the trail networks were most likely to optimize, we compared the observed ant networks to non-random networks generated by heuristic algorithms that optimized each objective independently (Methods). To compare observed networks with optimized networks, we performed two steps. First, we computed the percentile of observed networks with respect to random networks, and the percentile of optimized networks with respect to random networks. Typically, optimized networks had a percentile of 0%, but on some days, the percentile was slightly higher because the heuristic algorithm is not guaranteed to find the optimal network. Second, we compared the two percentiles for each objective to determine if the observed networks are more similar to random networks than to optimized networks.

When we examined the three pairwise correlations among average edge length, the total number of nodes, or average transition index, we found no correlations that were consistent in magnitude and direction in all four colonies (Table S1). By contrast, we found that the fourth objective (total edge length) was positively correlated with the total number of nodes, both when data from all 4 colonies were analyzed together (Spearman rank correlation, *R* = 0.62, *p* < 0.001) and separately (Spearman rank correlation, *R* > 0.70, *p* < 0.02 in all colonies). Thus, in our statistical tests, we did not compare total nodes and total edge length simultaneously. Instead, we examined whether trail networks reduce total edge length, and if so, whether they do so by using fewer nodes, or by using shorter edges. To do this, we first compared observed networks against random and optimized networks, to test whether trail networks significantly minimize total edge length. Then, to learn how the trail networks minimize total length, we examined whether networks minimized the total number of nodes, average edge length, or both.

### Turtle ant trail networks favor coherent trails over shortest edges

The ants’ networks optimized the maintenance of coherent trails. The observed networks were much more similar to the optimized networks in the extent to which they minimized the total number of nodes and transition index than average edge length (Fig 3A). The mean ± S.D. percentile differences between random and observed networks were: 47 ± 9% for total number of nodes; 42 ± 20% for average transition index; and 1 ± 44% for average length (n = 36 in all cases). The mean ±S.D. percentile differences between observed and optimized networks were: 3 ±8% for the total number of nodes; 6 ± 20% for average transition index; and 49 ± 44% for average length (n = 36 in all cases). We compared these two mean percentiles to determine whether observed networks were more similar to optimized than they were to random (Methods). Observed networks were more similar to optimized than to random networks for the total number of nodes (Wilcoxon signed ranks test, *n* = 36, *W* = 6, *p* < 0.001) and average transition index (Wilcoxon signed ranks test, *n* =36, *W* =76, *p* < 0.001), but the observed networks were more similar to random than to optimized networks for average edge length (Wilcoxon signed ranks test, *n* = 36, *W* = 180, *p* < 0.05).

**Fig. 3:**
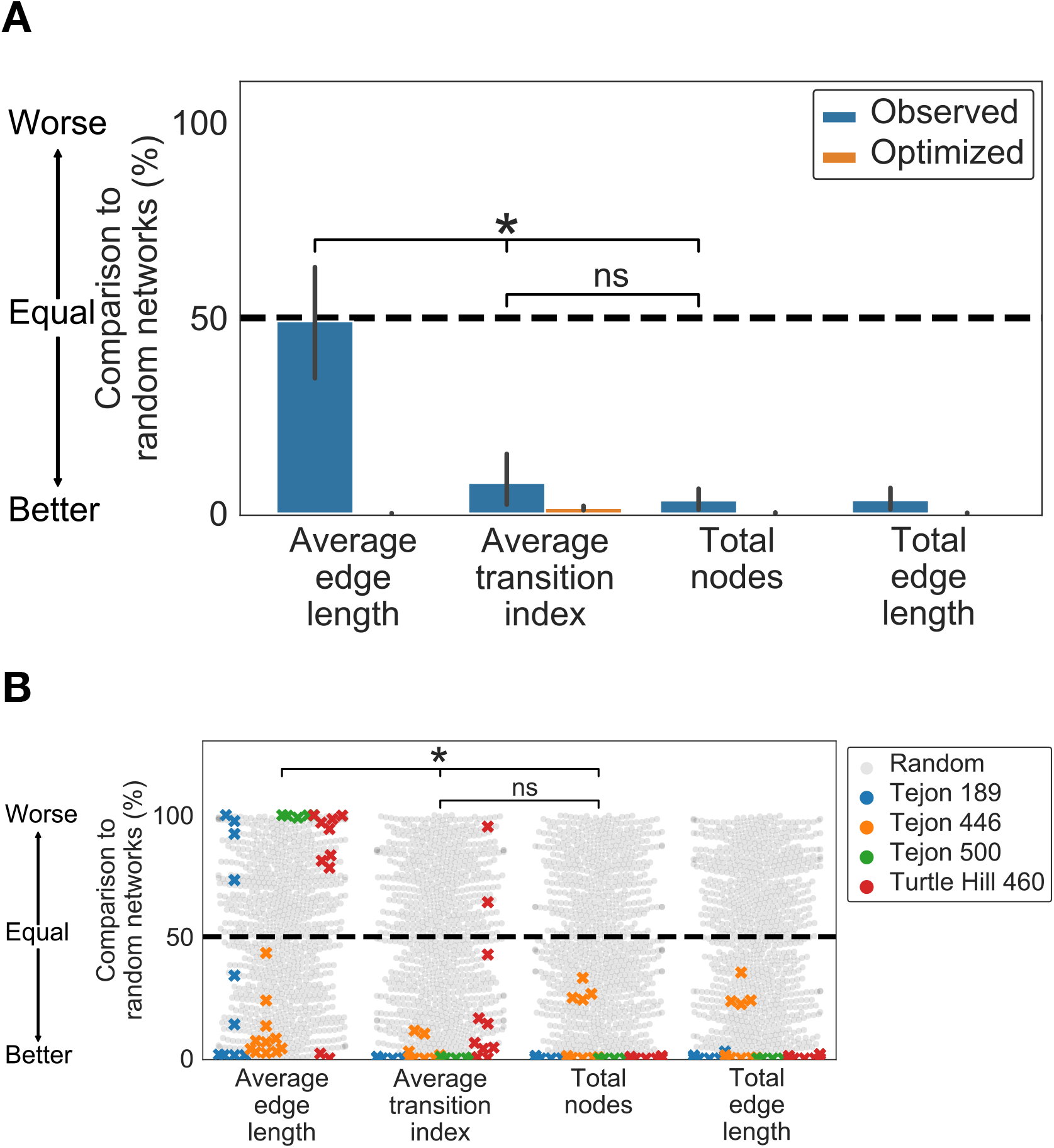
Comparison of random, optimized, and observed networks. A) Similarity of random, optimized, and observed networks measured as mean percentile (Methods). A percentile of 50 (dashed line) indicates that the observed or optimized network optimized the objective to the same extent as the average random network; the lower the percentile, the better the network optimized the objective compared to random networks. Error bars show standard errors of the mean of 36 non-independent networks. Asterisks represent p < 0.001, Conover test. B) A swarmplot of the percentile of each observed network, and the percentiles of a sample of the random networks. Gray circles represent the percentiles of random networks, and x’s represent the percentiles of observed trail networks, shown with a different color for each colony. Values below 50 indicate a network that performed better than the median random network for that objective, and values above 50 represent a network that performed worse. Asterisks represent *p* < 0.001, Conover test.

The ants’ networks also minimized total edge length significantly better than random networks did (Fig 3A–B). The mean ± S.D. percentile difference between random and observed networks was 46 ± 8%; the mean ± S.D. percentile difference between observed and optimized networks was 3 ± 8% (n = 36 in all cases). Observed networks were more similar to optimized than they were to random networks (Wilcoxon signed ranks test, *n* = 36, *W* = 4, *p* < 0.001). This indicates that turtle ant trail networks minimize total edge length. Because trail networks minimize the total number of nodes but not average edge length, we conclude that trail networks minimize total edge length by using fewer nodes rather than by using shorter edges.

When we compared how well the 3 uncorrelated objectives were optimized by the observed networks, relative to the day-to-day dependent random null model, there were significant differences among objectives (Friedman test; *Q* = 6.5; *p* < 0.05). When data for all colonies were analyzed together, trail networks minimized average transition index significantly more than they minimized the average edge length (Conover-Iman test [59, 60], *T* = 6.547 and 8.691, *p* < 0.001; Fig 3A–B) and minimized the total number of nodes. There were no significant differences between the extent to which observed networks, compared to random networks, minimized the average transition index and the total number of nodes (Conover test, *T* = 2.144, ns; Fig 3A–B).

Colonies did not differ overall in how well the observed networks met the 3 uncorrelated objectives, relative to the random networks (Friedman Test, *Q* = 2.2, ns), but there were differences among colonies in how well they optimized the objectives, apparently due to differences in the local vegetation in which the colonies traveled. Unlike the other 3 colonies, Turtle Hill 460 did not minimize average transition index significantly more than average length, relative to random networks. While on some days its trail networks minimized average transition index more than average length, on other days this did not occur, and the two objectives were not significantly different (Conover test, *T* = 2.135, ns). Unlike the other 3 colonies, Tejon 446 did not minimize the total number of nodes significantly more than it minimized average length (Conover test, *T* = 1.285, ns).

We also compared observed networks with random networks generated by the second null model in which random networks were generated independently for each day (Methods). The results were nearly identical: compared to random networks, the observed networks minimized the total number of nodes and average transition index, but did not minimize total length (Text S1,Fig S2)

### Day to day changes in trail networks

From day to day, progressive changes in the trail networks were more likely to optimize coherence than to minimize average edge length (Fig 4A–E). Fig 4A shows an example of a day-to-day change that minimized both the total number of nodes and average transition index. From day 9 to day 10, the network changed from the path shown in yellow to the path shown in blue, thus eliminating the 6 nodes circled, including 2 nodes with TI–2 and one node of TI–4, in favor of a path with nodes all of TI–1 (Fig 4A). Overall, trails in Tejon 189 (Fig 4B) consistently minimized the total number of nodes and average transition index but fluctuated in average edge length (days 4 to 9). In Turtle Hill 460 (Fig 4D), the network consistently minimized the total number of nodes, reduced TI significantly on days 4-10 compared to initially, on days 1-3, and fluctuated in average length (days 3 to 10). Changes in the nests and food sources used led to new networks with lower transition indices (days 2 to 10). There was an initial decrease in average edge length (days 4 to 7), due to a rupture on day 6 of a node leading to a 95cm edge (one of the longest edges we measured) rather than to a choice of shorter edges, and then the trails increased in average edge length (days 7 to 14). Overall, these changes suggest that from day to day, turtle ant networks are more likely to minimize transition index and reduce the number of nodes than to reduce the average edge length. This is consistent with previous observations of other colonies in which the number of nodes were reduced over time [51].

**Fig. 4:**
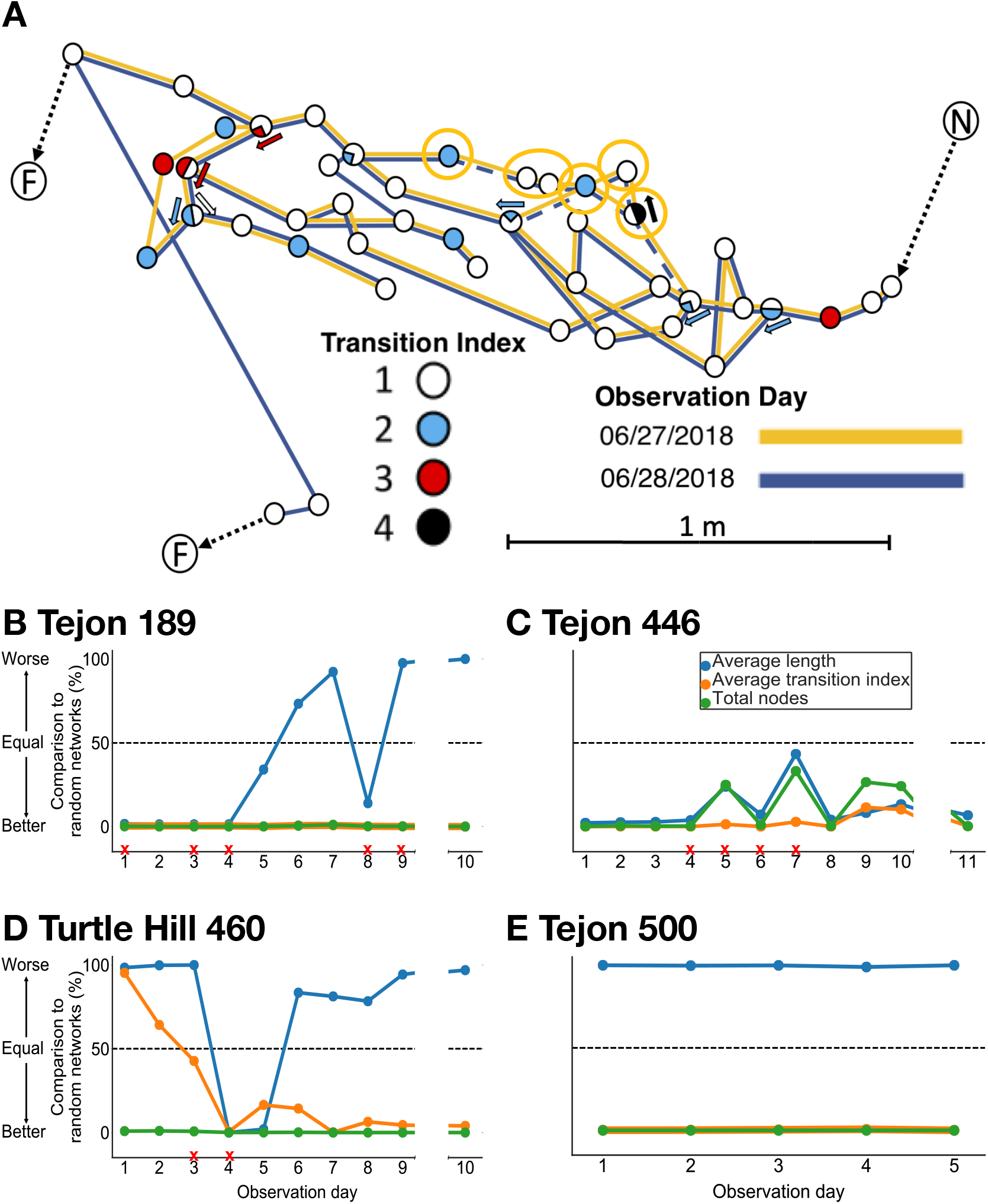
Comparison of random and observed networks over time. A) Part of the trail network of Tejon 189, showing a change from one day to the next. Symbols for transition index are the same as in Fig 2B: TI–1, open circles; TI–2, blue; TI–3, red; TI–4, black. Solid yellow lines show trails used on day 9; solid blue lines show trails used on day 10; yellow circles and dashed blue lines show trails linking the six nodes that were used on day 9 but not on day 10. For many nodes, not all edges are shown to simplify the illustration. In general nodes with higher TI have more edges. B–E) Day to day changes in mean percentile for each objective. A red ‘X’ indicates one or more ruptured edges on that day. A list of which dates correspond to which observation days in each colony is given in Methods. In B–D, the break in the x-axis between observation days represents an interval of about a month.

Using the day-to-day independent null model (Methods), we find the same trends over time; networks tended to minimize the number of nodes and average transition index rather than average edge length (Fig S2).

### Turtle ant networks form loops that promote coherence

Turtle ant networks form loops in their paths, consisting of small, temporary alternative paths with the same start and end points, and over time, all but one of these paths tends to be pruned away [51]. Loops are often considered to decrease the efficiency of routing networks [2, 3], but may increase robustness by offering alternative paths if links are broken. Here we hypothesize that loops may occur because trails tend to form along the most rapidly reinforced nodes, and a particular sequence of rapidly reinforced nodes may naturally form a cycle in the graph.

Compared to available loops in the vegetation with the same start and end points (Methods), the observed loops tended to use nodes with lower transition indices, and fewer nodes. We compared the centered ranks (Methods) of the average transition index, number of nodes, and average edge length, of paths connecting the same two start and end points on each trail in observed and available loops. The more negative the value of the centered rank, the lower the average transition index was on an observed path compared to all of the random paths within the available vegetation with the same start and end points. The mean ± S.D. of the centered rank in observed loops for average transition index was −2.13 ± 4.13; for number of nodes was −2.51 ± 4.09 and for average edge length was 0.60 ± 2.95; in all cases, significantly different from 0 (T-Test, |*T*| > 3.2, *p* < 0.01). These results suggest that loop formation in trails may occur as a consequence of selecting trails that have lower transition indices.

## Discussion

The trail networks of arboreal ants link nests and find temporary food sources in changing vegetation. They respond to dynamic environments by maintaining the coherence of their trails. Minimizing the transition index and minimizing the number of nodes contribute to coherence because they both diminish the risk of losing ants from the trail. Like water flowing over a rocky stream bed, turtle ants tend to use trails most conducive to the flow of ants. When the physical configuration of a node provides a trajectory that successive ants are likely to follow, the chances are reduced that turtle ants wander off the path and lay pheromone trails that can lead other ants also to leave the trail.

Trail networks must also balance the tradeoff between exploration and coherence. Exploration is necessary for the colony to construct trails [61, 30, 42, 17, 41, 39], search for new resources [62, 63, 64, 65, 66, 40, 54] and repair breaks [51, 52, 34], and the connectivity of the vegetation (Table 1) influences the probability that ants that leave the trail will return to another node on the trail. There appears to be a small probability of exploration at each node [51, 52], so the probability of leaving the trail accumulates with more nodes traversed. Thus, minimizing the number of nodes traversed reduces opportunities for the ants to get lost.

Nodes with transition indices of 1 keep the trail on the same plant [51]. The vines and trees of the tropical dry forest tend to have long internode distances to reach the sunlight at the edge of the canopy [67]. By staying on the same plant, ants are led to resources at the edge of the canopy, such as flowers that provide nectar.

Minimizing distance traveled is often considered the main objective in engineered routing algorithms and ant colony optimization [24, 22, 23], since rapid communication is often needed between any two nodes in the network. We showed that the trail networks of turtle ants minimize total edge length, a measure of distance travelled, by minimizing the total number of nodes, rather than by minimizing the average edge length. We also show that turtle ant trail networks minimize the average transition index, which is an NP-complete optimization problem (Methods). Minimizing the total number of nodes and minimizing average transition index both promote coherence by increasing the probability that ants stay attached to the trail. It seems that keeping ants together on the trail, thus maintaining the routing backbone for collecting and distributing resources, is more important than minimizing the lengths of edges used. In previous work [52], we proposed a model for the algorithm used to maintain and repair networks. This algorithm did not distinguish between minimizing the number of nodes and the distance traveled, since each edge had equal length. Our results here show that further work is needed to develop an algorithm that captures the role of physical variation in the environment.

The costs of additional nodes include the loss of ants from the trail, and the pursuit of fruitless paths, which may detract from the colony’s ability to distribute resources among its many nests, and make fewer ants available to recruit effectively when a new food source is discovered. In addition, each node provides opportunities for encounters with other, competing species traveling in the same vegetation [68, 69, 70, 58]. Our results here suggest that, as in engineered networks [71, 72], the cost of including a node in the turtle ant network may vary among nodes, because whether other ants follow an exploring ant that leaves the trail at that node depends on the node’s physical configuration. We assigned transition indices here by visual inspection of the vegetation, and we hope in future work to investigate further the effects of node configuration on path formation using biophysical models based on 3D scanning data.

This study of 4 colonies across about 10 days, as well as previous studies of another 21 colonies [50, 51] shows that in general, colonies choose similar trajectories through the vegetation, but there are also interesting differences. Further work is needed to examine variation among colonies [73] in the extent to which they maximize the objectives we studied here, how such variation corresponds to differences among sites in vegetation, and whether variation in colony behavior is associated with colony reproductive success.

We compared observed ant networks with two kinds of random networks, first with random networks that modified the previous day’s trail, and second with random networks generated from scratch each day, sampled broadly from the space of all possible networks. Both models gave similar results: both performed significantly worse than the ant networks for minimizing the total number of nodes and average TI. This suggests that the ant networks are very far from being random, and that there are an extremely large number of sub-optimal networks in the vegetation. Moreover, the ant networks, using only local cues and de-centralized decision-making, are very close to performing as well as heuristically optimized algorithms that use centralized, global information.

Are there useful applications of the principle that variation in the environment provides useful constraints on routing network design? Evolved algorithms operating in the natural world can use structure in the environment to enhance coordination among distributed agents [74]. In engineering, taking into account the physical structure of the environment may improve the design of routing algorithms [75, 76, 77, 78, 79], for example, by reducing the search space of possible routing paths and steering network construction away from parts of the terrain that are difficult to reach. This could be beneficial in applications such as robot swarms, where distributed agents must coordinate in complex environments using a communication backbone to explore new terrain, exploit of temporary resources, and maintain security against intruders [58].

## Materials and Methods

### Observation of trail networks

We observed trail networks in 4 colonies, for a total of 36 (10 + 11 + 10 + 5) non-independent trail networks. The trail networks of four turtle ant colonies (Turtle Hill 460, Tejon 446, Tejon 189, and Tejon 500) were observed between 06/20/2018 and 06/28/2019 at La Estación Biológica de Chamela in Jalisco, Mexico. Tejon 189 was observed 10 times: Once per day from 06/22/2018–06/28/2018, and once per day on 07/01/2018, 07/04/2018, and 08/03/2018. Tejon 446 was observed 11 times: once per day from 06/20/2018–06/22/2018, once per day from 06/24/2018–06/28/2018, and once per day on 07/01/2018, 07/04/2018, and 08/03/2018. Turtle Hill 460 was observed 10 times: twice on 06/21/2018, once on 06/22/2018, once per day from 06/24/2018–06/26/2018, and once per day on 06/28/2018, 07/02/2018, 07/05/2018, and 08/03/2018. Tejon 500 was observed 5 times: once per day from 06/24/2019–06/28/2019. In observations made in August 2018, we did not find changes in the surrounding vegetation mapped in July. Not every network was observed on each day. Colonies were found as in previous work [50, 51]. For this study, we chose colonies that had a large part of their trail networks low enough in the canopy that all vegetation within 5 nodes of the path used was accessible for mapping and measuring. The colonies appeared to be of about the same size, based on observations in comparison with previous work in which size was estimated with mark-recapture measures [50], but here we did not measure colony size.

Networks were mapped each day between 09:00 and 13:00 using the same methods as in previous work [51], using visual inspection of the paths the ants took through the vegetation at a rate of at least 10 ants per 15 min in one direction. Previous work showed this to be the approximate threshold for a trail that would be used consistently throughout the day (e.g., Fig. 7 in Gordon (2017) [51]); lower rates of flow are associated with brief temporary searches that do not lead to an established trail, presumably because the pheromone trail, with an estimated exponential decay rate of 0.02 units per second [52], is not sufficiently reinforced. This rate of flow was used to give each edge a binary assignment of either “used” or “not used”. No emigration events were observed. Turtle ants usually stop foraging in response to the presence of other ant species [50, 51]; it is rare to see an ant of another species on an active trail, and none were observed during the course of our study. To track changes in the path and terminals used each day, each node was assigned a number, and enough nodes were marked, with small labels or stickers on a dead end branch near the node, to identify all the same nodes the next day.

The assignment of the transition index to the vegetation at each node was made using the following features of the vegetation:

- TI–1 (Fig 1A): a node linking two edges on the same plant.
- TI–2 (Fig 1B): a node that links one plant to another along a trajectory through a node that is likely to be the same for successive ants.
- TI–3 (Fig 1C): a node that links one plant to another plant with more than one possible trajectory through the node.
- TI–4 (Fig 1D): a node that links one plant to another with many possible trajectories that is often changed by conditions such as the wind.

On the first day of observation, we mapped the network used by the ants on that day as well as most of the surrounding vegetation up to 5 nodes away. When new nodes were added to the paths used by the ants, we then mapped most of the available vegetation up to 5 nodes from the new nodes. When nodes were deleted from the vegetation, they were also deleted from the networks used in simulations. All nodes had at least 3 edges, including the one along which the ant approached the node, and thus there were at least 2 choices of edges to take in the direction it was traveling. In each day of observation, we recorded which edges and nodes were used by the ants. As observed previously [51], each colony’s trail network changed which nodes it used from day to day, although each day’s path conserved some parts of the previous day’s. The terminals used differed from one day to another, as food sources are found and abandoned, new nests are added, and an existing nest can be lost when the branch it is in breaks [50, 51]. On some days, for unknown reasons, some nests were not connected to the network (Fig S1). In the course of the observations reported here, the path of the ants never went outside the range of the 5 nodes around the trail that were originally mapped. The number of nodes observed in each network is shown in Table 1.

### Assigning directionality to edges

Previous experiments with marked ants show that ants tend to use particular routes from a nest [50], and tend not to turn around on the trail. To account for this, for all edges we defined a direction relative to one terminal; the outbound direction went away from it, and the inbound direction went towards it. We restricted the analysis to paths that proceed in the outbound direction.

### Modeling the transition index as a line graph

The transition index of a node is an estimate of the probability that successive ants will take the same trajectory from an edge (*u, v*) through node *v* to edge (*v, w*). Thus a transition index involving node *v* is assigned to a particular incoming edge used to reach *v* and a particular edge used to leave *v*.

To model the transition index, we used the line graph of *G*, where *L_G_* = (*V_L_, E_L_*). The nodes *V_L_* = *E_G_*, and the edges *E_L_* = {((*x, y*), (*y, z*))|(*x, y*), (*y, z*) ∈ *E_G_*}. The line graph creates a node for each edge in *G* and connects two nodes if they correspond to two adjacent edges in *G*. Every edge in the line graph denotes a transition in the original graph; traversing the edge ((*u,v*), (*v, w*)) ∈ *L_G_* corresponds to starting at *u*, going to *v*, crossing the junction at *v*, and then going to *w*. We assigned every edge in *L* a transition index. Fig 5 illustrates this process.

**Fig. 5:**
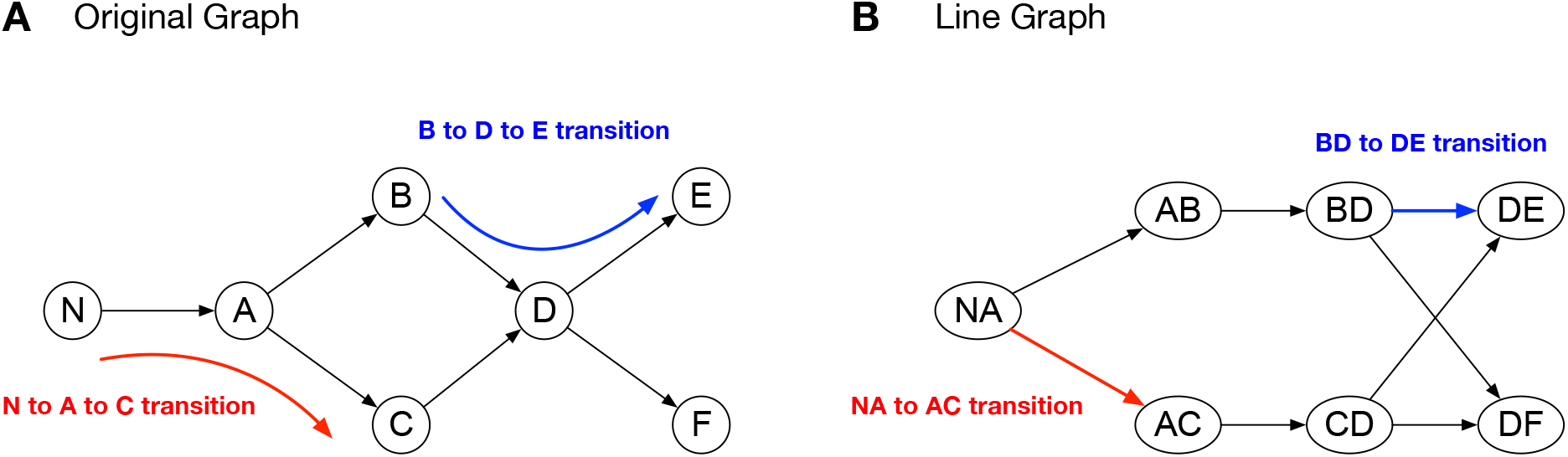
Converting the network to the line graph. A) Original graph. Nodes correspond to junctions in the vegetation, and edges correspond to links, such as a branch or stem, between junctions. B) Line graph. Every edge in the original graph has a corresponding node in the line graph. Two nodes are connected in the line graph if they correspond to two adjacent edges in the original graph. Transitions in the original graph used in this example are highlighted in red and blue.

### Generating random trail networks

We compared the average edge length, total number of nodes, and average transition index in observed and random trail networks. The random networks were simulated in the graph made from the map of the paths used by ants and the surrounding vegetation, using the number of nodes, edge lengths and transition indices measured in the vegetation. The random trail networks may include loops, as did the observed networks.

We used two methods to generate random trail networks. The first null model, called the *day-to-day dependent model*, starts with an initial random network, and then randomly modifies it from day to day to connect the same terminals, nests, and food sources that were used by the turtle ants. Results are reported in the text. The second null model, called the *day-to-day independent model*, generates a new random network from scratch every day. Results are reported in Text S1.

For the day-to-day dependent model, we start by creating a random network that connects all of the terminals (nests and food sources) observed to be used on the first observation day. This initial network is different for each simulation. To generate a random network for each subsequent day, we modify the previous day’s trail network to reflect any changes to the set of terminals used by the turtle ants. If a new terminal, such as a new nest or food source, is included in the observed network on a given day, we add a path going from that terminal to the random network for that day. If a terminal, such as a nest or food source, is not used on a given day in the observed network, we remove the path going from the terminal to the random network for that day.

Formally, we start by generating an initial network *G*_1_ for the first observation day:

1. **Input:** Graph *G* = (*V_G_, E_G_*), terminals *X* ⊆ *V_G_*.
2. Create an empty graph *G*_1_; add a random terminal *x* ∈ *X* to *G*_1_.
3. Choose a random terminal *x*′ ∈ *X* that has not already been added to *G*_1_.
4. Perform a random walk on *G* that starts at *x*′ and stops when it touches any node *n* in *G*_1_.
5. Add all edges and nodes touched by the random walk to *G*_1_.
6. Repeat steps 3–5 until all terminals have been added to *G*_1_.

Let *G_t_* be the trail network generated randomly for day *t*. We now describe the process of constructing network *G*_*t*+1_ for day *t* + 1. First, we define the following term: when we add a terminal *x*′ to the trail network, we perform a random walk that starts at *x*′ and stops when it touches a node *e* that is already in the network; we call *e* an *endpoint*.

To construct *G*_*t*+1_ we use the following procedure:

1. For each terminal *x_i_* that was used on day *t* and not used on day *t* + 1, let *W_i_* be the random walk through the vegetation that started at *x_i_* and was originally used to connect *x_i_* to the existing network (either on day *t* or an earlier day). Let *e* be the first endpoint that appears in *W_i_*.
2. Remove all nodes in *W_i_* that are never touched after *e* is touched for the first time.
3. For each terminal *x_j_* that was used on day *t* + 1 and not on day *t*: perform a random walk that starts at *x_j_* and stops when it touches any node *u* in *G*_*t*+1_. Add all nodes and edges touched by the random walk to *G*_*t*+1_.

After randomly generating an initial network *G*_1_ for day *t* = 1, we use the above procedure to generate a network for day 2; and then continue this process for all observation days. This procedure reflects the observation that that turtle ant trails use similar paths from day to day. We ran 100,000 simulations in which we generated a different initial network.

In Text S1, we present results from the second null model in which each random network is generated independently of the random network from the previous day. On each day, we use the same 6 step procedure that was used to create an initial network, to create a new network from scratch using the same terminals as the observed network.

### Approximating optimal networks

We evaluated networks generated by heuristic algorithms designed to optimize each objective. Finding the network that optimizes total nodes is a variant of the classic Steiner tree problem (an NP-complete problem) [71, 72], and so we applied a standard approximation algorithm [80]. To optimize average length, we used the following algorithm:

1. **Input:** Graph *G* = (*V_G_, E_G_*) and terminals *X* ⊆ *V_G_*
2. Create an empty graph *T*.
3. Add edges *T* in order of increasing length until *T* contains all terminals *X*, and all terminals are in the same connected component.
4. Remove all nodes and edges not part of the same connected component as the terminals *X*
5. Remove edges from *T* in decreasing order of length; only remove edges that decrease the average length without disconnecting *T*

To optimize average transition index, we use a nearly identical algorithm, but we add and remove edges in order of transition index rather than length. To optimize average transition index, we use a nearly identical algorithm, but we add and remove edges in order of transition index rather than length. To optimize total edge length, we used a directed Steiner tree approximation algorithm devoped by Charikar, et. al. [80].

### Comparing observed and random networks

For each of the 4 colonies (Tejon 189, Tejon 446, Turtle Hill 460, and Tejon 500), and for each objective (average length, total nodes, and average transition index, total edge length), we computed the similarity of each observed network on a given day to 100,000 random networks that connected the same set of terminals, using Equation (1).

To compare the observed and random networks, we computed for each network the following four objectives: 1) Average edge length: the average length of all edges in the network; 2) Total number of nodes: the total number of nodes in the network; and 3) Average transition index: the average transition index over all transitions in the network; 4) Total edge length: the total length of all edges in the network.

For each objective, we computed a value of *φ*_ants_ for the observed network and *φ*_1_, *φ*_2_,…, *φ_n_* for the *n* random networks. We measured the similarity between the observed network and the random networks for that objective as the percentage of random networks that have a lower value for the objective:

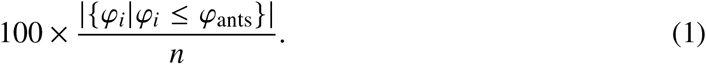

An observed network attains the same value as the median random network when its percentile is 50%. The closer the percentile is to 0, the better the observed network optimized the objective, and the closer the percentile is to 100, the better the random network optimized the objective. This percentile-based approach is unit-less, making it possible to compare performance for objectives that differ in the range of values.

#### Statistical comparison of observed and random networks

To compare how well each of the three uncorrelated objectives were optimized, and to compare how well each of the four colonies optimized objectives, we used the non-parametric Friedman test that makes no assumptions about the distribution of the data [81]. We considered how networks from each of the 4 colonies optimize 3 uncorrelated objective measures for a total of 4 × 3 = 12 comparisons. To avoid pseudoreplication in day to day measurements, for each objective and each colony we computed the average of the percentiles over observation days. We found the average value for each objective and each colony, and performed two Friedman chi-square tests: one testing for differences among colonies, and one testing for differences among objectives. We then compared the objectives using post hoc Conover-Iman non-parametric tests [59, 60], with a Bonferroni correction for multiple comparisons and a significance threshold of *p* = 0.05.

### Comparing observed and optimized networks

We would expect observed networks not to perform as well as optimized networks on every objective, since optimized networks are found by an algorithm that draws on information about the entire map, whereas the ants make decisions based on only local information. Thus, rather than comparing observed and optimized networks directly, we tested whether observed networks were closer to optimized networks than would be expected by chance. To compare how similar observed and random networks were to optimized networks, we first computed the percentile of observed networks with respect to random networks, and the percentile of optimized networks with respect to random networks. We then tested if the observed networks were significantly more similar to random networks than to optimized networks. For each objective, we evaluated the difference between the observed network and random network as: 0.50 – (the observed network’s percentile), and the difference between the observed network and the optimized network as: (the observed network’s percentile) – (the optimized network’s percentile). We found this difference for each of the 36 observed networks and each of the 4 objectives. The 36 networks were formed by 4 colonies and are snapshots of 4 dynamic networks over time, rather than 36 fully independent networks. We then used the 36 differences for each objective to test whether the observed values were closer to the optimal value for that objective than to the median random value, with a Wilcoxon signed ranks test. We recognize that the 36 differences were not independent because they include sets of observations from 4 colonies.

### Loops

We compared the average transition index and number of nodes in observed loops with loops that were available in the surrounding vegetation. A loop was defined as two or more outbound paths that start and end at the same source and target nodes. For each observed loop, we computed all possible paths in the vegetation between the corresponding source and target nodes. We ranked all paths based on the average transition index of the path, and compared this to the rank of the average transition index of the paths used by the ants. We repeated the same measure for total number of nodes and average edge length.

We computed the *centered rank* of each path used by the ants as follows: We ranked *n* available paths in the vegetation, by the average transition index of nodes in the path, total number of nodes in the path, or average length of edges in the path. The ranks of the paths are 1, 2, 3,… *n*, and the median rank is *r_m_* = (*n* + 1)/2. For each observed path, we computed its centered rank by subtracting the median rank from that path’s rank. Using average transition index as an example, the centered rank is 0 when the observed loop had the same average transition index as the median random loop, negative when the observed loop has a lower average transition index, and positive when the observed loop has a higher average transition index. We performed a similar analysis for the total number of nodes and average edge length.

### Connectivity

To calculate the connectivity of the vegetation (Table 1), we estimated how many nodes are required for an ant that leaves the trail to return to the trail. For each edge (*u, v*) connecting a node on the trail to a node off the trail, we found the length, in number of nodes, of the path with fewest nodes from *v* back to a node on the trail. We measured connectivity as the smallest number of nodes in a path back to the trail, averaged over all edges (*u, v*) leading off the trail on all days.

### Optimizing transition index is NP-complete

Here we show that the problem of constructing the trail network that connects a given set of terminals while minimizing the average transition index of the network is NP-complete. To do this, we start by showing that finding the minimum average-weight path between two vertices in a graph is NP-complete. In this problem, we are given a graph *G* = (*V_G_, E_G_*) and two vertices *u, v* ∈ *V*. The goal is to find a path 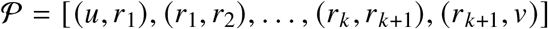 that minimizes the average edge length: 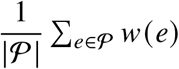, where *w*(*e*) defines the length of edge *e*. This problem differs from the classic shortest path problem, which seeks a path with minimal total edge length.

The standard method for considering the complexity class of an optimization problem is to consider the equivalent decision version of the problem: given a graph *G* = (*V_G_, E_G_*), two vertices *u, v* ∈ *V_G_*, and an integer *k*, we ask: is there a path from *u* to *v* whose average weight is ≤ *k*?

#### Lemma 1.

*Finding the minimum average-weight path is NP-Complete*.

*Proof*. First, we show that this problem is in the class NP. If we are given a path from *u* to *v*, we can verify that the path is a valid *u*-*v* path, and that the average edge weight is ≤ *k*. This certificate will clearly be of polynomial length, and we can verify that it is correct in polynomial time.

Next, we show that the problem is NP-hard. We proceed via reduction from the Hamiltonian path problem, which is NP-Complete. Given a directed graph *G* = (*V_G_, E_G_*), the directed Hamiltonian path problem seeks a path that touches every vertex *v* ∈ *V_G_* exactly once. We construct a graph *G*′ as follows: *G*′ contains all of the vertices in *G* along with two additional vertices, *s* and *t*. We assign a weight of 1 to all of the original edges in *G*. We add a directed edge from *s* to every vertex *v* ∈ *V_G_*, and we add a directed edge from every vertex *v* ∈ *V_G_* to *t*. We assign a weight of 2 to all edges that include either *s* or *t*.

The smallest possible average edge weight for any path from *s* to *t* is 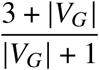, and such a path exists if and only if *G* has a directed Hamiltonian path.

The reduction requires adding 2 new nodes, and 2|*V*| new edges to *G*. Thus the reduction clearly takes only polynomial time.

We now use Lemma 1 to show that optimizing average transition index is NP-Complete. Let *L* = (*V_L_, E_L_*) be a graph whose edge weights correspond to transition indices, which we represent using the line graph (Fig 5). Given a set of terminals *X* ⊆ *V_L_*, we seek to find a subtree *F_X_* ⊆ *L* that minimizes the average value of all edge weights in *F_X_*. We require that *F_X_* be connected without cycles to represent the fact that cycles in turtle ant trails are typically pruned.

Once again we consider the decision problem: given a line network *L* = (*V_L_, E_L_*), a set of terminals *X*, and an an integer *k*, does *L* contain a subtree *F_X_* whose average weight is ≤ *k*?

#### Lemma 2.

*Finding the subtree F_X_ that optimizes average transition index is NP-Complete*.

*Proof*. First, we prove that the problem is in NP. Given *V_L_, X, k*, if we are presented with a subgraph *F_X_* as a certificate, we can verify that *F_X_* is a valid solution. We check that *F_X_* is valid subgraph, that *F_X_* is connected, that *F_X_* contains every terminal node *X*, and that the average weight of the edges in *F* is ≤ *k*. Clearly the size of *F_X_* is polynomially bounded, and we can verify that *F_X_* is a valid solution in polynomial time.

To show that the problem is NP-hard, we proceed via reduction from the minimum average-weight path problem above. Given *G* and vertices *u* and *v*, we simply treat *G* as the line graph (*L*), and designate terminals *X* = {*u, v*}; this means *F_X_* is simply a *u*-*v* path. We seek the trail that optimizes the average transition index between terminals *u* and *v*. By construction, this is the minimum average-weight path from *u* to *v* in the line graph, meaning there is a subgraph *F_X_* with average weight of ≤ *k* if and only if the original graph has a *u*-*v* path with average edge weight ≤ *k*. Further, the reduction clearly takes only polynomial time.

The classic Steiner tree problem, in which the objective is to minimize *total* length, is known to be fixed-parameter tractable in the number of terminals [82]; that is, if the number of terminals is held constant then the problem can be solved in polynomial time. By contrast, optimizing transition index is not fixed-parameter tractable for the number of terminals; our reduction shows that optimizing average transition index is NP-hard even if we restrict the number of terminals to 2. Thus, optimizing average transition index is more complex than even the classic NP-complete Steiner tree problem.

## Acknowledgements

We thank Ibai Zabaleta and especially Taggert Butterfield for assistance in the field; and Daniel Beck, Katharine Renton, and the staff at the Estación Biológica de Chamela for their help. We are grateful for the comments of anonymous reviewers of a previous version of the manuscript.

## Author Contributions

AC: data curation, formal analysis, methodology, software, validation, visualization, writing (original draft preparation). JARM: conceptualization, funding acquisition, methodology, project administration, software, supervision, visualization, writing (original draft preparation). CA: data curation, investigation, visualization. SN: conceptualization, funding acquisition, methodology, project administration, supervision, writing (original draft preparation). DMG: conceptualization, data curation, funding acquisition, investigation, methodology, project administration, supervision, visualization, writing (original draft preparation).

## Data Availability Statement

All data collected and analyzed will be archived at Stanford Digital Repository at the Stanford Libraries. Python code is available at: https://github.com/DiODeProject/TurtleAntObjectives.

---

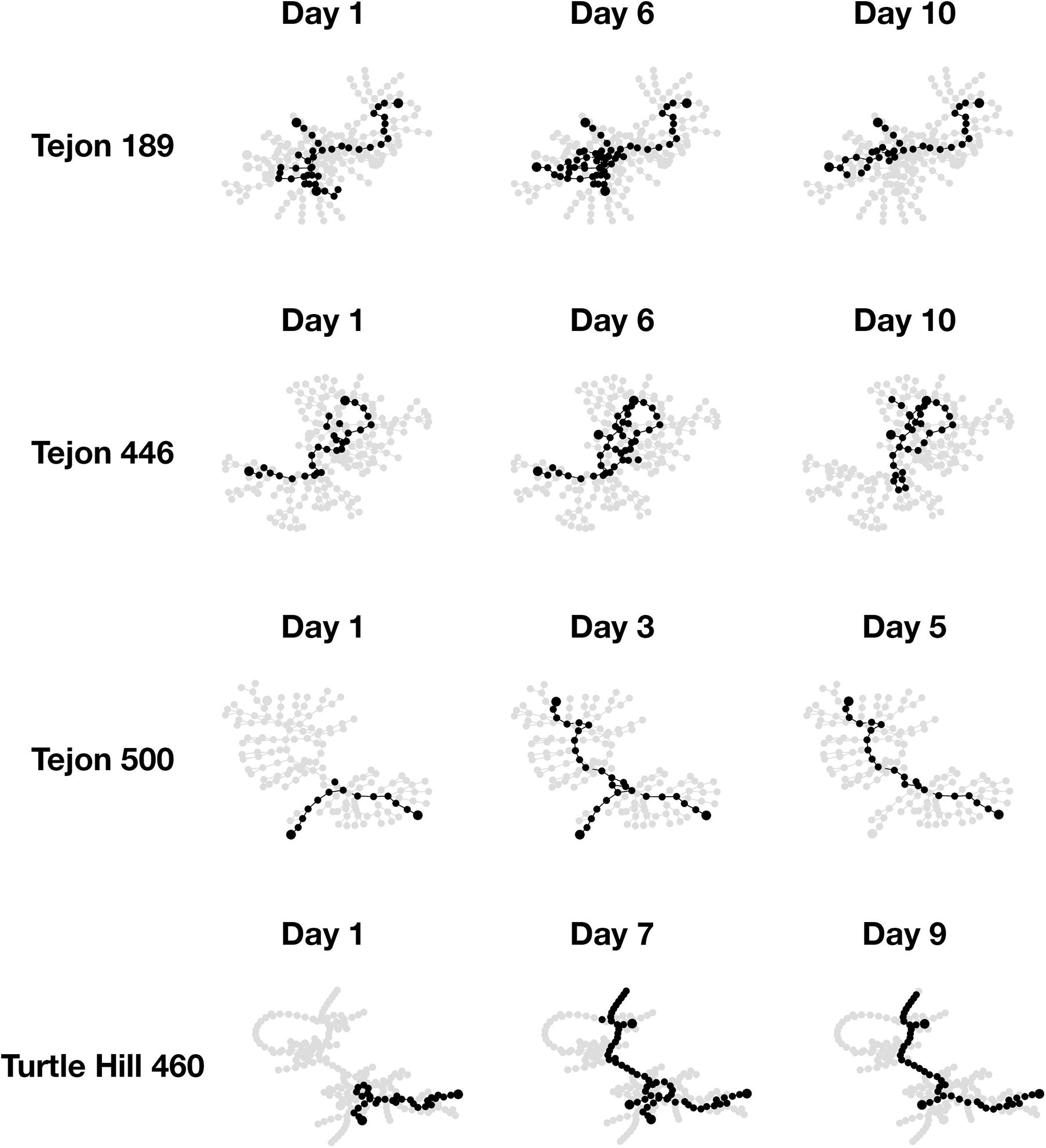

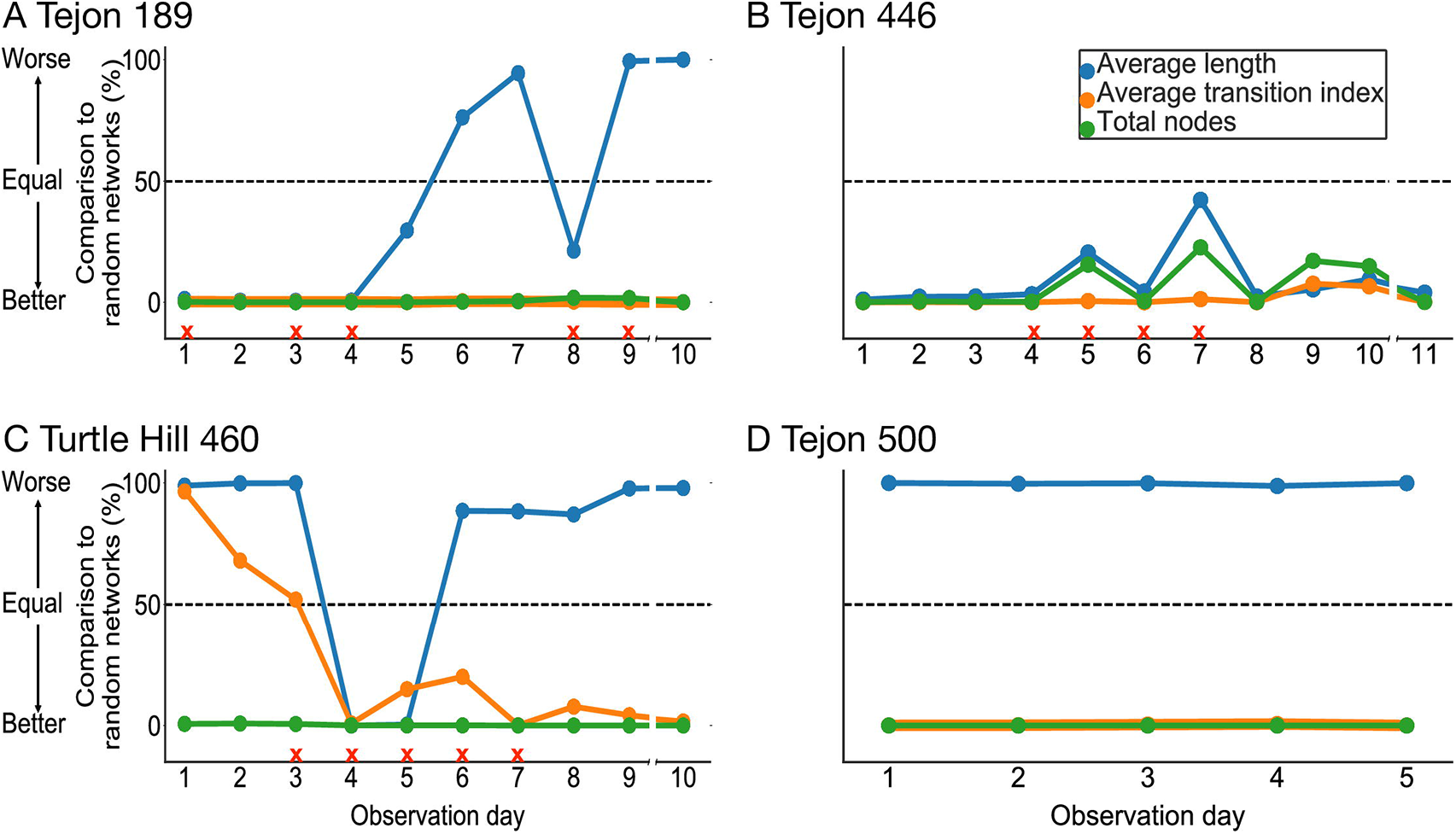

