## supplementary-material for "Better tired than lost: turtle ant trail networks favor coherence over short edges"

### Pairwise correlations between candidate objectives objectives

| Objective 1 | Objective 2 | Tejon 189 | Tejon 446 | Tejon 500 | Turtle Hill 460 |
|---|---|---|---|---|---|
| Average transition index | Total nodes | 0.41 | 0.10 | 0.97* | -0.51 |
| Average transition index | Average edge length | -0.37 | 0.03 | -0.99* | 0.58 |
| Total nodes | Average edge length | 0.24 | 0.25 | -0.97* | -0.92* |

Table S1: **Pairwise correlations between candidate objectives.** We computed Spearman's rank correlation between each pair of candidate objectives. Asterisks indicate $p < 0.01$.

### Changes in the trail networks over time

### Comparing trail networks to an alternate null model

The main text presents the results of the day-to-day dependent null model in which the structure of each random trail network depended on the structure of the trail network generated on the previous day. Here we report the results of a second null model comparing the observed trail networks with trail networks that were generated independently from one another as follows:

1. **Input:** Graph $G = (V_G, E_G)$, terminals $X \subseteq V_G$.

2. Create an empty graph $R$; add a random terminal $x \in X$ to $R$.

3. Choose a random terminal $x' \in X$ that has not already been added to $R$.

4. Perform a random walk on $G$ that starts at $x'$ and stops when it touches any node in $R$.

5. Add all edges and nodes touched by the random walk to $R$.

6. Repeat steps 3–5 until all terminals have been added to $R$.

We evaluated how well the observed trails optimize the three objectives using the day-to-day independent model. The results are nearly identical (Figure S2) to those for the day-to-day dependent model. First, when we compared how well the 3 objectives were optimized by the observed networks, relative to the day-to-day independent random null model, there were

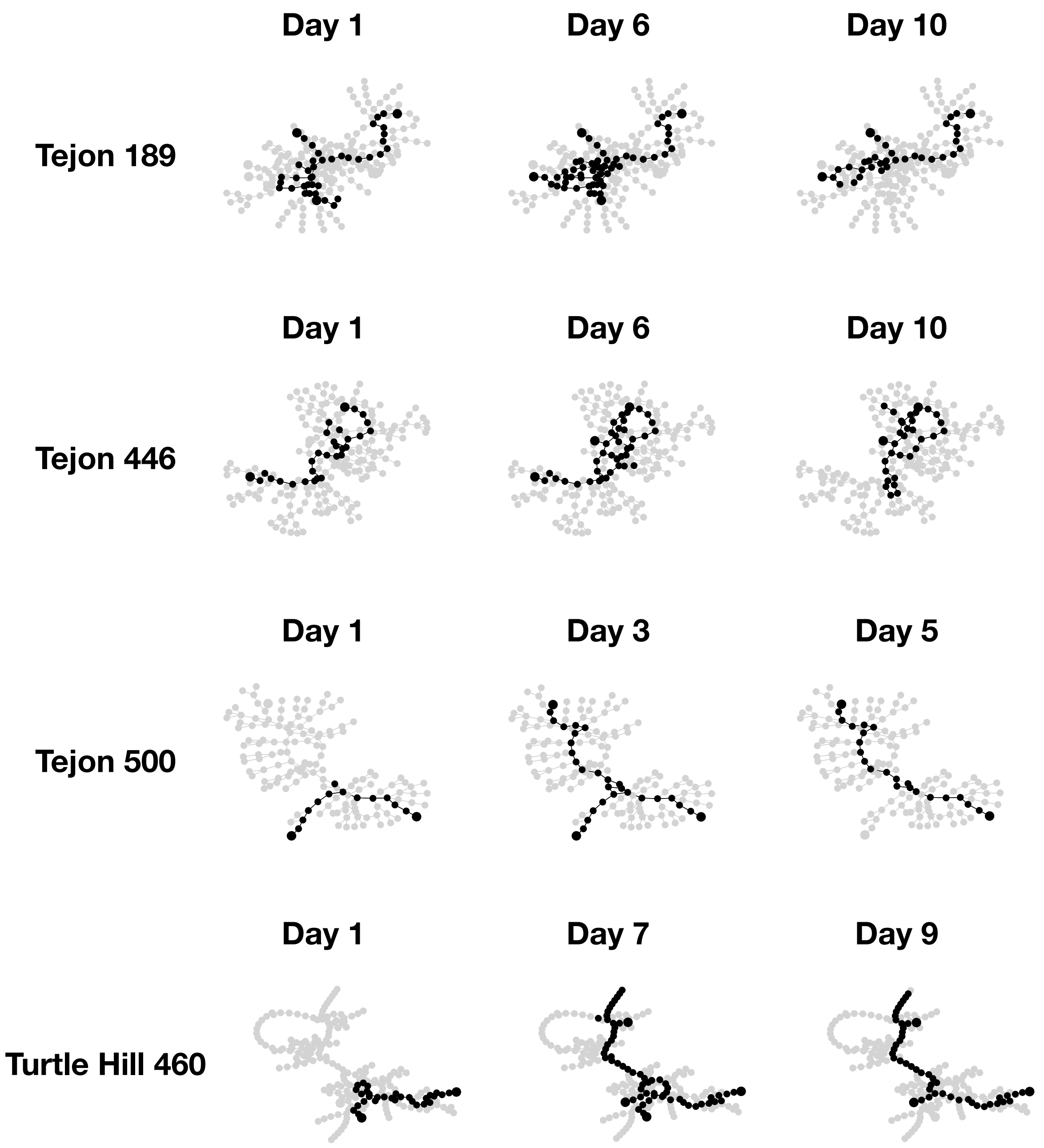


Fig. S1: **Changes in trail networks over time.** For each colony, the trail network is shown for the first, middle and last day of observation. Black dots represent nodes in the trajectory taken by the ants on that day, and gray dots represent nodes in the surrounding vegetation that were not used on that day.

significant differences among objectives (Friedman test; $Q = 6$; $p < 0.05$). Trail networks minimized average transition index (Conover test , $T = 6.181$, $p < 0.001$) and minimized the total number of nodes (Conover test, $T = 8.312$, $p < 0.001$), significantly more than the distance traveled. There were no significant differences between the extent to which observed networks, compared to random ones, minimized the average transition index and the total number of nodes (Conover test, $T = 2.130$, ns).

The comparison of observed versus optimized networks using the day-to-day independent model showed that observed networks were significantly closer to optimized networks for total nodes and average transition index than for total length.

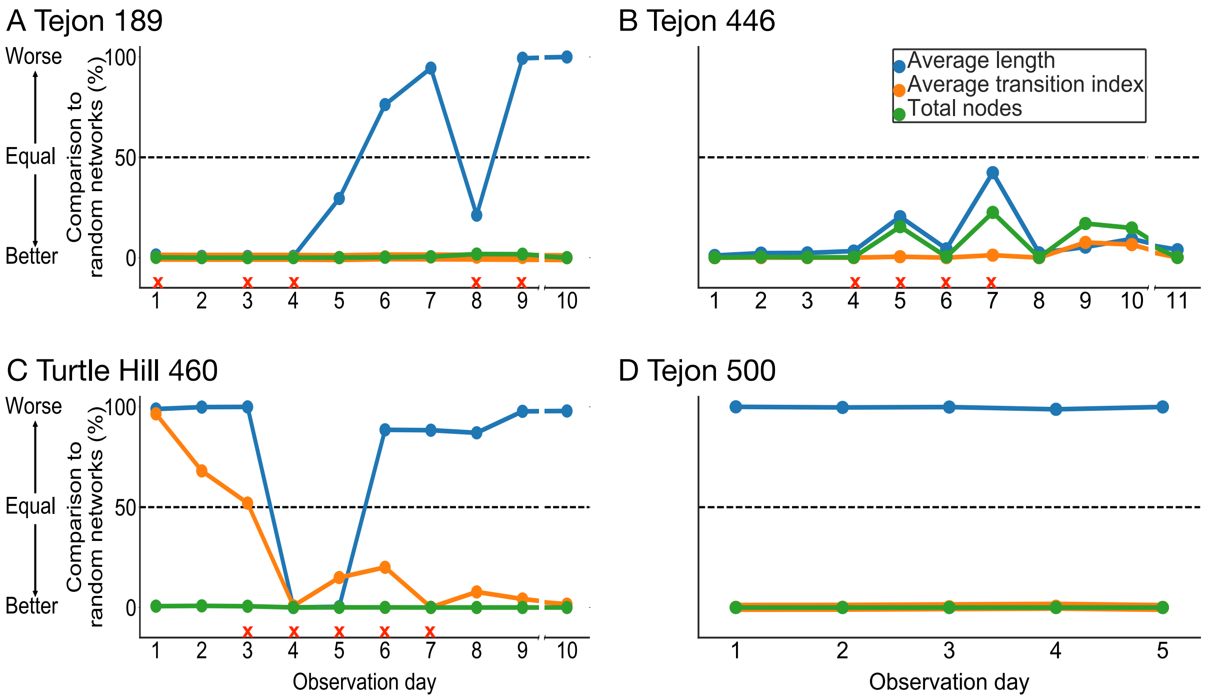


Fig. S2: **Comparison of random and observed networks using the day-to-day independent model.** Day to day changes in mean percentile for each objective. A red 'X' indicates one or more ruptured edges on that day. A list of which dates correspond to which observation days in each colony is given in Methods. In C–E, the break in the x-axis between observations days represents an interval of about a month.
